# Minimal protein list for chromatin replication-coupled symmetric histone recycling revealed through *in vitro* reconstitution

**DOI:** 10.1101/2024.11.18.624094

**Authors:** Fritz Nagae, Shizuko Endo, Yasuto Murayama, Tsuyoshi Terakawa

**Affiliations:** Department of Biophysics, Graduate School of Science, Kyoto University, Kyoto, Japan; Department of Chromosome Science, National Institute of Genetics, Shizuoka, Japan; Department of Genetics, Graduate University for Advanced Studies (SOKENDAI), Shizuoka, Japan

## Abstract

Upon eukaryotic DNA replication, symmetric histone recycling from parental to daughter strands is vital for transmitting epigenetic information to the next generations. Recent genome-wide sequencing studies have identified several protein regions contributing to symmetric histone recycling in a cell. However, a minimal list of proteins for symmetric histone recycling remains unknown. Here, we successfully reconstituted histone recycling with ∼30 purified proteins and analyzed the products digested by Micrococcal nuclease with the newly developed pipeline called a Repli-pore-seq in which nanopore sequencing and deep-learning-based classification were combined. As a result, we confirmed that the histones were recycled symmetrically to the lagging and leading strands. The recycled histones form tetrasomes or hexasomes and are deposited on the DNA sequences energetically favorable for forming nucleosomes. We also observed the discordance of the recycled position between lagging and leading strands on the GC-rich DNA sequences. Among proteins dispensable for chromatin replication, the lagging-strand maturation factors, Fen1/Cdc9, were also dispensable for symmetric recycling. The removal of Pol δ disrupted the recycling symmetry, and that of Ctf4 or Csm3/Tof1 altered the recycling position. These findings provide critical insights into the molecular players and mechanisms underlying symmetric histone recycling.

## INTRODUTION

Epigenetic information, which is encoded in chromatin, is crucial to regulate transcriptional patterns and to determine functions and fates of cells^1–8^. Faithful inheritance of epigenetic information across generations is essential for maintaining cellular identities. The nucleosome, a basic unit of chromatin, is a protein/DNA complex in which 147 base-pairs (bp) DNA wraps around a histone octamer composed of one H3/H4 tetramer and two H2A/H2B dimers^9^. Post-translational modifications (PTMs) in histones are one representative carrier of epigenetic information to regulate gene expressions^4,6,10^. Recycling of modified histones from parental chromatin to either of two replicated DNA strands is the first step of epigenetic inheritance, which is reportedly coupled with DNA replication^1–8,11^. Symmetric recycling of parental histones to the replicated strands is vital for maintaining cellular identity after cell division^12–15^.

Deep-sequencing-based studies using eSPAN^16^ and SCAR-seq^17^ have demonstrated that parental histones are symmetrically recycled to the lagging and leading strands at a replication fork. The techniques have identified protein regions to regulate symmetric histone recycling^16–26^. Mcm2, a subunit of a replicative helicase, has a histone-binding domain on its N-terminal tail and contributes to recycling to the lagging strand^17^. Additionally, other subunits of replisome (Ctf4^18^, Pol α^18,19^, Pol δ^19,20^, PCNA^20^, and Tof1^21^) are proposed to participate in the recycling to the lagging strand. On the other hand, Dpb3 and Dpb4, non-essential subunits of Pol ε, are essential for recycling to the leading strand^16^. Mrc1, one subunit of the replisome, is involved in histone recycling to both lagging and leading strands^22,23^. A histone chaperone FACT captures parental histones ahead of a replication fork and contributes to recycling to both strands^24,25^. A deep-sequencing study for cells with combinatorial mutations in Mcm2, Dpb4, and Pob3 (a subunit of FACT) demonstrated that defects of histone recycling to daughter strands are additive^25^, indicating the coexistence of multiple histone-recycling pathways. Together, these experiments using cells suggested that the symmetry of histone recycling relies on balance among the recycling pathways.

Although these genome-wide sequencing studies have revealed symmetric histone recycling, how the replisome itself contributes to the symmetry has been elusive. In cellular environments, coexisting enzymes such as transcription, repair, and nucleosome remodeling machinery potentially modulate replication progression and nucleosome formation and positioning^27–30^. Moreover, certain PTMs, which have been used as markers of old histones, can be added to newly synthesized histones by histone-modifying enzymes^6,10^. These enzymes, which alter the epigenetic states of nucleosomes, complicate the interpretation of the genome-wide sequencing studies. Furthermore, combinatorial depletion of proteins disrupted cell growth and downstream analyses, making it a challenge to identify the minimal protein list for symmetric histone recycling. Therefore, *in vitro* reconstitution of chromatin replication, which allows adding proteins to or removing those from the system and isolating the contribution to histone recycling, is highly desired.

*In vitro* reconstitution of biological systems is a powerful tool for understanding the fundamental mechanisms of complex cellular processes. The reconstitution of eukaryotic DNA replication identified the minimal protein list of DNA replication and molecular mechanisms of the process^31–33^. In addition, a combination of the *in vitro* DNA replication and other reconstitutions (*e.g.*, nucleosome^34–39^, R-loop^40^, or cohesin^41^) have successfully revealed the coordination of eukaryotic DNA replications and the other processes. These studies demonstrated that the reconstituted replisome can recycle histones onto the replicated DNA strands in the absence or presence of histone chaperones^34,42^. However, the gel-electrophoresis-based assays used in the studies did not provide critical information regarding the recycling ratio to the lagging and leading strands and recycling position, which has inhibited the elucidation of the molecular players and mechanisms underlying symmetric histone recycling.

Here, we reconstituted histone recycling with ∼30 purified proteins and analyzed the products digested by Micrococcal nuclease with the newly developed pipeline called Repli-pore-seq in which nanopore sequencing and deep-learning-based classification were combined. The results confirmed that histones are symmetrically recycled to the lagging and leading strands, forming tetrasomes or hexasomes on DNA sequences energetically favorable for nucleosome formation. Interestingly, we identified positional discordance of recycled histones between the lagging and leading strands on GC-rich DNA regions, though they have the same DNA sequence. Furthermore, we found that the lagging-strand maturation factors, Fen1/Cdc9, are not essential for symmetric and faithful recycling, suggesting that maturation may happen after recycling. Although our gel-electrophoresis-based assays demonstrated that Pol δ, Ctf4, and Csm3/Tof1 are dispensable for histone deposition to the replicated DNA, the removal of Pol δ disrupted the recycling symmetry and the removal of Ctf4 or Csm3/Tof1 caused alterations in recycling positions. These findings shed light on the molecular components and mechanisms driving symmetric histone recycling, paving the way for a deeper understanding of epigenetic transmissions.

## RESULTS

### In vitro reconstitution of chromatin replication-coupled histone recycling

We mixed purified proteins for *Saccharomyces cerevisiae* DNA replication and histone chaperone FACT with chromatinized DNA substrates to reconstitute *in vitro* chromatin replication, as described previously^33,34,38^. We assembled nucleosomes onto a 4.1 kilo-bp (kbp) plasmid DNA with histones, Nap1, and ISW1a. Then, we exchanged buffer using gel-filtration column (Fig. 1A). Almost all the free histones were filtered out by the column while histones bound to DNA were eluted (Supplementary Fig. 1A lanes 5 and 7). We added ORC, Cdc6, and Mcm2-7/Cdt1 to assemble MCM double hexamer on the nucleosome substrate (Fig. 1A). After MCM loading to ARS consensus sequence (ACS) site in the plasmid, we initiated DNA replication by adding firing factors (DDK, S-CDK, Sld2, Sld3/7, Dpb11, Cdc45, GINS, Csm3/Tof1, Ctf4, Mrc1, RFC, PCNA, RPA, Pol α, Pol δ, Pol ε, and Mcm10) and histone chaperone FACT in the reaction (Fig. 1A). Newly replicated DNA was labeled by incorporation of biotin-dUTP in reaction mixture (Fig. 1A). We incubated the reaction for 20 minutes to complete duplication of the DNA substrates (Fig. 1B lane 2, 1C & Supplementary Fig. 1B). We found that FACT was indispensable for replicating chromatin (Fig. 1B lane 1, 1C & Supplementary Fig. 1B), consistent with the previous biochemical experiments^34,41^.

**Figure 1:**
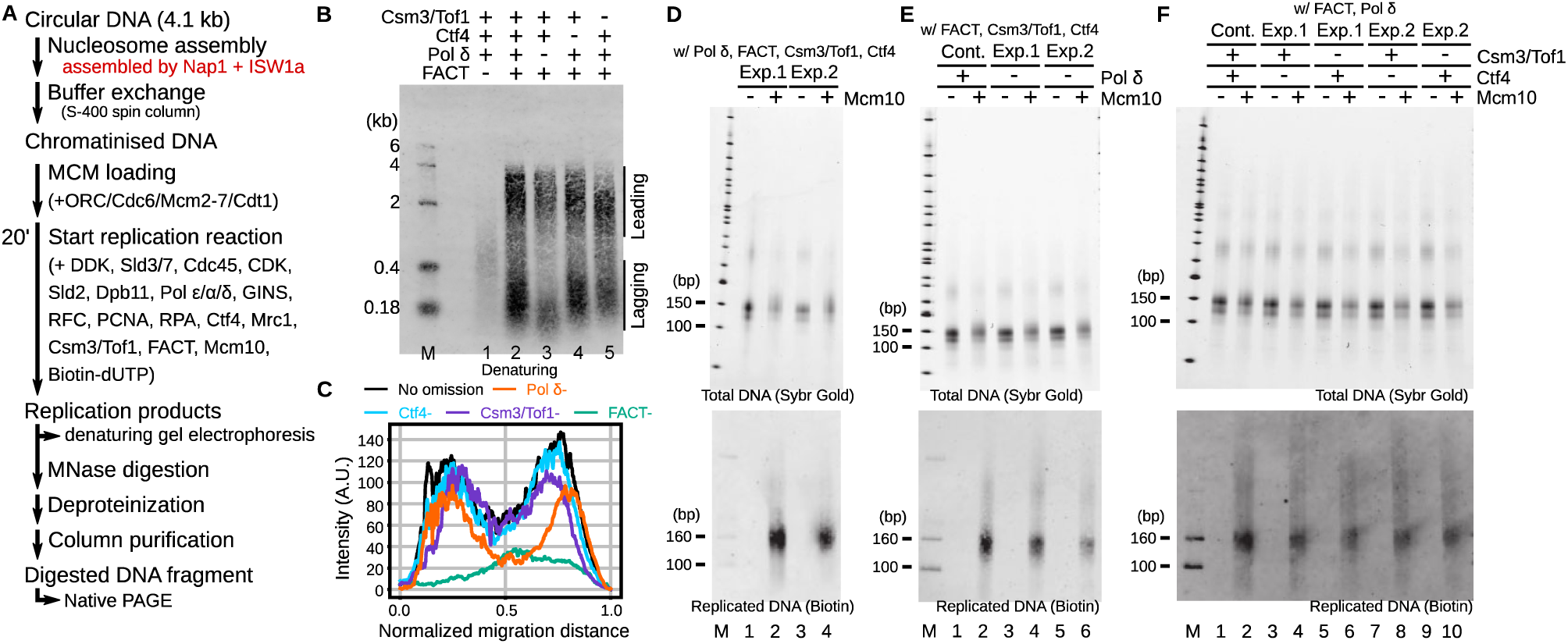
*In vitro* reconstitutions of chromatin replication. (A) The protocol of *in vitro* DNA replication with plasmid DNA forming nucleosomes. (B) The alkaline denaturing agarose gel image of replicated DNA before MNase digestion. ‘M’ denotes a marker. (C) Lane profiles of replicated DNA in (B). (D– F) The native polyacrylamide gel images of total (top) and replicated DNA (bottom) after MNase digestion. The replication reactions of (D) were conducted according to the protocol in (A). The reactions of (E) and (F) were conducted by removing Pol δ and Ctf4 or Csm3/Tof1 in the firing step in (A). ‘M’ denotes a marker.

Previous studies have demonstrated that salt gradient dialysis (SGD) assembles irregularly spaced nucleosome arrays while Nap1 and ISW1a (NI) assemble regularly spaced and phased nucleosome arrays to generate nucleosome-depleted-region (NDR) on the ACS site^35,36,43^. To test which of the SGD- or NI-chromatin is suitable substrate for the *in vitro* chromatin replication, we assembled nucleosome onto the plasmid DNA using SGD and repeated the replication assay (Supplementary Fig. 1C). The replication reaction on the SGD chromatin completed when we assembled nucleosome by mixing histones with the plasmid DNA at an equimolar ratio (Supplementary Fig. 1C). However, slight or no signal of the replicated strand was detected when the amount of histone is increased to twice that of the plasmid DNA (Supplementary Fig. 1C). This indicated that NI-chromatin is a suitable substrate for efficient chromatin replication, consistent with the previous study^35^. Therefore, we used the NI-chromatin for all the experiments hereafter.

After replication reactions, we digested the products with Micrococcal nuclease (MNase) to examine whether histones were deposited onto DNA (Fig. 1A). We observed ∼150 bp DNA fragments in the reaction without Mcm10, replication initiation factor^44,45^, when staining all the DNA strands, supporting successful nucleosome formation on parental DNA (Fig. 1D lanes 1 and 3 of the top panel). We also found the ∼150 bp DNA band in the reaction containing Mcm10 (Fig. 1D lanes 2 and 4 of the bottom panel) when detecting only replicated and biotinylated DNA, while no signal was detected in the absence of Mcm10 (Fig. 1D lanes 1 and 3 of the bottom panel). The ∼150 bp biotinylated DNA band did not appear when we used naked DNA as a replication substrate (Supplementary Fig. 1D). These data supported that the purified proteins listed in Fig. 1A are sufficient for successful recycling of histones onto newly replicated DNA, consistent with the previous results^34,42^.

Next, we omitted proteins not essential for chromatin replication and examined the effect of removing a factor on histone recycling. Previous studies have demonstrated that Pol δ, Ctf4, and Csm3/Tof1 are dispensable for DNA replication^33,38^. Pol δ is responsible for the lagging strand synthesis^46,47^, which Pol ε can replace. Ctf4 and Csm3/Tof1 are at the front edge of the replisome and work as scaffolds for other replication proteins (e.g., Pol α and FACT)^18,21,37,48,49^. We repeated the replication assays without Pol δ, Ctf4, or Csm3/Tof1 (Pol δ−, Ctf4−, or Csm3/Tof1−). The replicated products in the Ctf4− and Csm3/Tof1− reactions were similar with their presence (Fig. 1B lanes 4 and 5, 1C & Supplementary Fig. 1B). On the other hand, removing Pol δ reduced the length and amount of the lagging strand while effects on the leading strand were moderate (Fig.1B lane 3, 1C & Supplementary Fig. 1B), consistent with the results using naked DNA as a substrate^33^. After the MNase digestion, we observed ∼150 bp biotinylated DNA bands in the same way as their presence (Fig. 1E lanes 2, 4, and 6 & 1F lanes 2, 4, 6, 8, and 10). These results indicated that Pol δ, Ctf4, and Csm3/Tof1 are dispensable for histone deposition to replicated DNA, although removing Pol δ shows moderate effects on chromatin replication.

### Deep learning model to detect biotinylated DNA with nanopore sequencing

To analyze the recycling ratio to the lagging and leading strands and recycling position, we established a pipeline called Repli-pore-seq in which nanopore sequencing and deep-learning-based classification were combined. In this pipeline, we first performed *in vitro* chromatin replication assays as shown above (Fig. 1A, 2A & 2B). After the MNase digestion, DNA fragments are a mixture of template unmodified DNA and replicated biotinylated DNA since all substrates could not always be replicated (Fig. 2B). Immunoprecipitation-based purification of biotinylated DNA did not work due to the low efficiency of the reconstituted histone recycling. To computationally rather than biochemically “purify” the replicated DNA, we performed nanopore sequencing of the mixture with the MinION device (Oxford Nanopore Technologies) and classified the template unmodified DNA and the replicated biotinylated DNA based on their electric current signals (Fig. 2A to E). In the nanopore sequencing, one single-stranded DNA strand passes through a nanopore embedded in the membrane, and its sequence is identified based on changes in the current signal through occluding the pore by the nucleotides (Fig. 2C & 2D). Previous studies have revealed that nucleotide analogs show unique current patterns distinguishable from four canonical nucleotides^50,51^. We developed a deep-learning-based classifier to detect the current signals unique to the incorporated biotin-dUTP.

**Figure 2:**
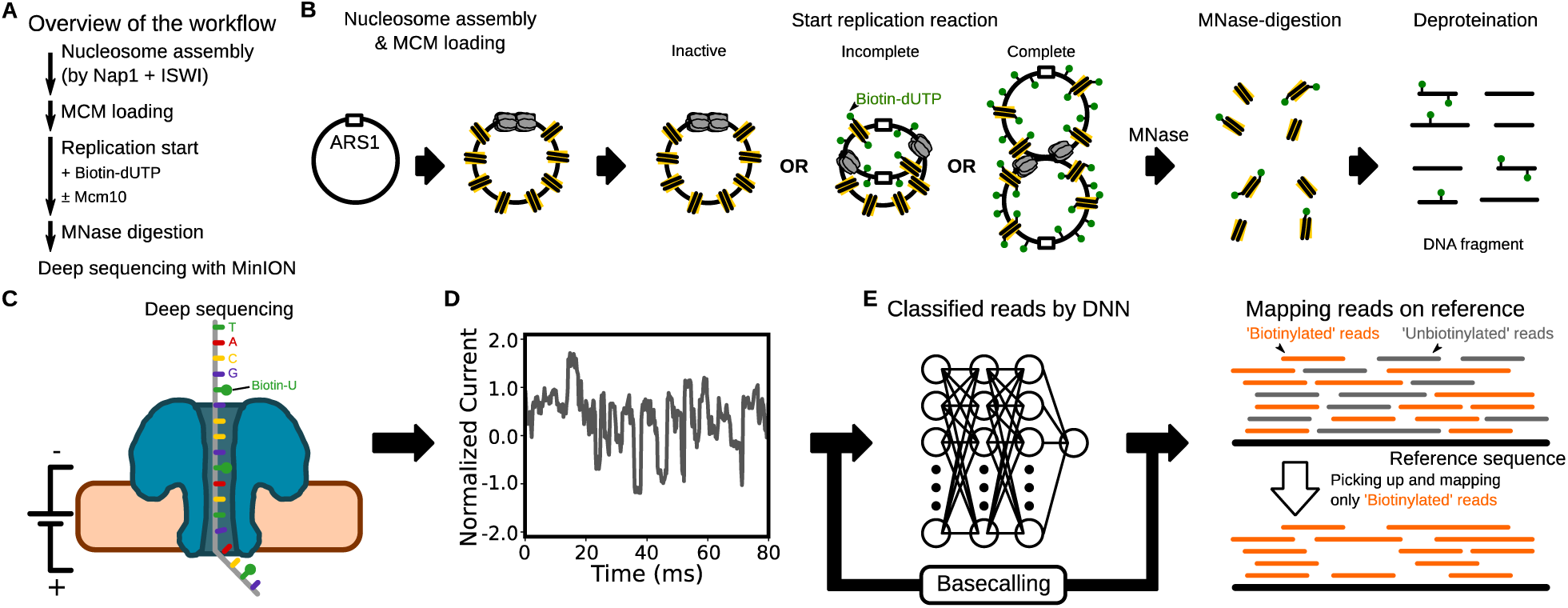
The Repli-pore-seq pipeline. (A) The protocol of the pipeline. (B) Schematic of *In vitro* chromatin replication, MNase digestion, and deproteination. (C) Schematic of nanopore sequencing using the MinION. (D) A representative raw signal obtained by MinION. The signals were segmented and converted to an array of the segments’ means, standard deviations, and durations. (E) Schematic of the deep-neural network model. The model is composed of residual blocks with convolution and LSTM layers. The values in the arrays obtained in (D) were converted into a scholar within the range [0, 1] and were fed into the network. The reads were labeled as ‘biotinylated’ if the output was ≥0.5; otherwise, they were labeled as ‘unbiotinylated’.

We combined the residual neural network with LSTM to establish the classifier (Fig. 2E, Supplementary Fig. 2A & 2B; See Methods for details). For training, we prepared biotinylated and unbiotinylated DNA substrates by PCR and measured the current signals of each substrate using the MinION device. The current signal was segmented according to its stepwise change, and each segments’ mean, standard deviation, and duration were calculated (Fig. 2D). Each series of the segments from unbiotinylated and biotinylated DNA was labeled as ‘0’ and ‘1’ respectively, and an equal number of the signals from each class were mixed to create the training data. We trained the classifier to predict whether each read contains biotin-dUTP nucleotides.

We evaluated the performance of the trained classifier. Predicted labels of unbiotinylated and biotinylated DNA substrates for the validation dataset were distributed around 0 and 1, respectively (Supplementary Fig. 2C). Few instances in both substrates showed the predicted labels around 0.5. Therefore, we considered the label below 0.5 as ‘unbiotinylated’ and the others as ‘biotinylated’. A confusion matrix showed that the accuracy and precision of the model were 98% and 99%, respectively (Supplementary Fig. 2C inset).

### Repli-pore-seq confirmed reconstituted symmetric histone recycling

We applied Repli-pore-seq to our *in vitro* reconstitution system of histone recycling. We found that 11.6% and 0.5% of the reads were classified as ‘biotinylated’ in the reaction with and without Mcm10 (Mcm10+ and Mcm10−, respectively) (Fig. 3A). The increased proportion of ‘biotinylated’ reads in the Mcm10+ reaction was consistent with the biotinylated DNA fragments detected on the gel (Fig. 1D). On the other hand, biotinylated DNA fragments were not detected on the gel in the Mcm10− reaction (Fig. 1D). Therefore, we considered the 0.5% ‘biotinylated’ reads as misclassified unbiotinylated DNA fragments. Based on this assumption, we estimated the fraction of genuinely biotinylated reads in the Mcm10+ reaction (See Methods for details). Hereafter, we consider that all the reads in the Mcm10− reaction correspond to parental DNA fragments and that the reads classified as genuinely biotinylated in the Mcm10+ reaction correspond to daughter DNA fragments.

**Figure 3:**
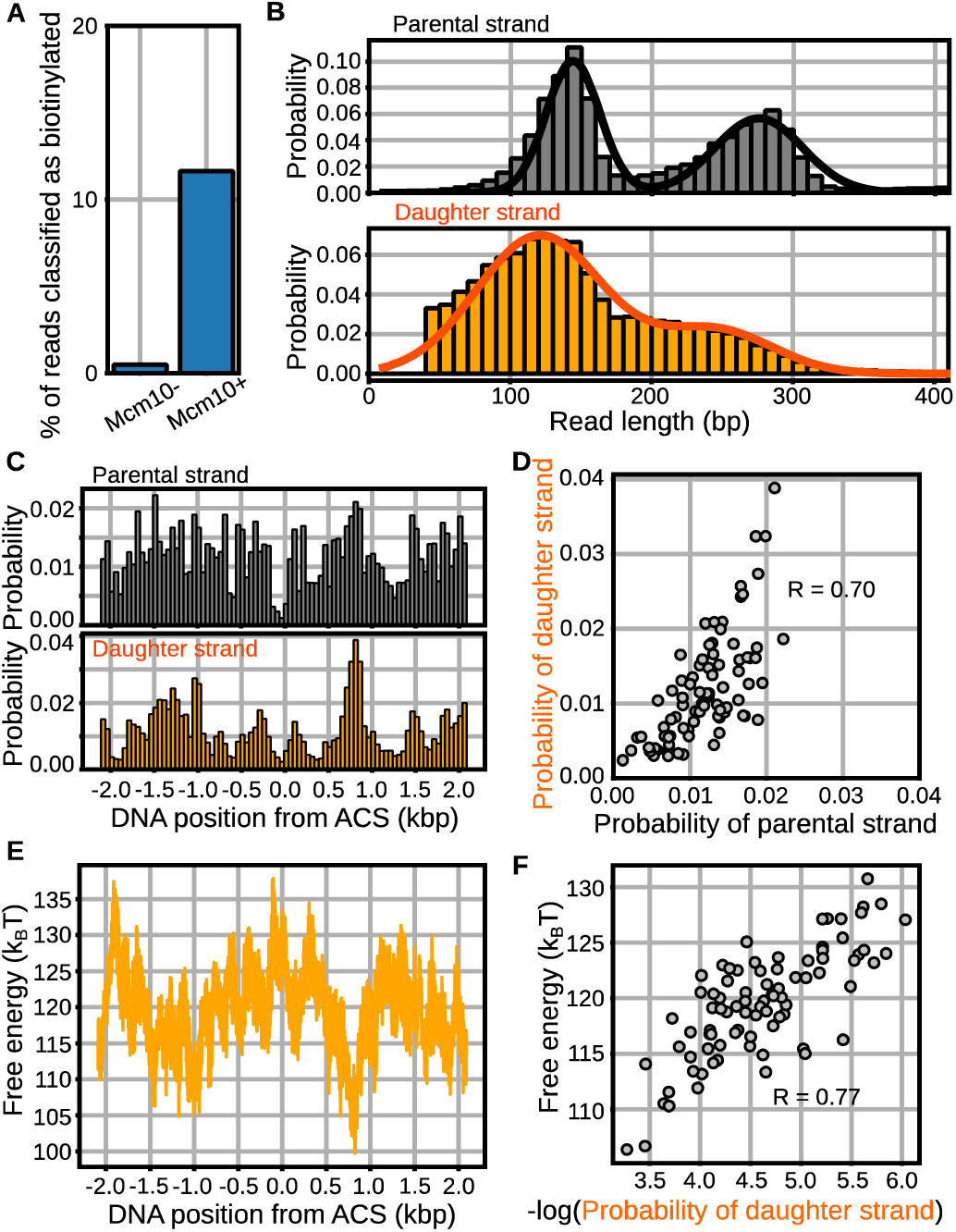
Repli-pore-seq of the reconstituted chromatin replication. (A) Fractions of biotinylated reads in the Mcm10− and Mcm10+ reactions. (B) The read length distributions of parental DNA fragments (top, gray) and daughter DNA fragments (bottom, orange). The distributions were fitted using two Gaussians. (C) The position distributions of histones on the parental DNA (top; gray) and the daughter DNA (bottom; orange). (D) The correlation plot for the position distributions of histones on the parental and daughter DNA in (C). (E) The theoretically calculated energy landscape of nucleosome formation along the template DNA. (F) The correlation plot for the energy landscape in (E) and the logarithm of the position distribution of histones on the daughter DNA.

To get insights into the oligomeric state of the recycled histones, we compared the read length of the parental DNA fragments with that of the daughter DNA fragments. We found two peaks in the read length distribution of the parental DNA fragments, which correspond to mono- (144 ± 19 bp) and di-nucleosome (275 ± 31 bp) (Fig. 3B & Supplementary Fig. 3A). The peak read length of daughter DNA fragments was 121 ± 45 bp, shorter than that of parental DNA fragments (Fig. 3B & Supplementary Fig. 3A). Also, the fractions ranging from 40 to 120 bp increased for the daughter DNA fragments (Fig. 3B & Supplementary Fig. 3A). These results suggested that some nucleosomes lost one or two H2A/H2B dimers to form hexasomes or tetrasomes after histone recycling.

Next, we compared the histone positions on the parental and daughter DNA. We used 200 bp or shorter reads for the analysis and defined a midpoint of the read as a position of histones. The position distributions of the parental and daughter DNA fragments were not uniform, but the patterns were reproducible for repetition (R = 0.98 and 0.99, respectively) (Fig. 3C & Supplementary Fig. 3B to D). On the other hand, the position distribution was almost uniform when we sequenced the naked plasmid DNA sheared by sonication (Supplementary Fig. 2D), showing that the read bias was negligible. The less frequent localization of histones around the ACS site supported the successful loading of replication initiation complex to chromatinized DNA (Fig. 3C). The position distribution of the daughter DNA fragments was moderately correlated with that of the parental DNA fragments (R = 0.70) (Fig. 3D), indicating that histone positions were conserved before and after recycling. Then, we theoretically calculated the free energy landscape of nucleosome formation from the DNA sequence^52^ and compared it with the logarithm of the position distribution of the daughter DNA fragments (Fig. 3E), finding that they are also moderately correlated (R = 0.77) (Fig. 3F). These results suggested that the histones are positioned on the DNA sequences energetically favorable for forming nucleosomes after recycling.

Because nanopore sequencing reads single-stranded DNA, we can classify reads of daughter DNA fragments into the lagging and leading strands according to their relative positions from the ACS site (Fig. 4A). 53.2% and 52.7% of the daughter DNA fragments were classified as the leading strand fragments in two repeated experiments (Fig. 4B), strongly suggesting that histones are symmetrically recycled to the lagging and leading strands in our *in vitro* reconstitution system, consistent with the observation in cells^16–26^. The read length distributions of the lagging and leading strand fragments were not significantly different (Fig. 4C & Supplementary Fig. 4A). Interestingly, while correlation between the position distributions from the repeated experiments was high (R = 0.99) (Supplementary Fig. 4B & 4C), that between the lagging and leading strand fragments were moderate (R = 0.52) (Fig. 4D). In two repeated experiments, we reproducibly observed that the recycled positions between lagging and leading strands were roughly similar but not entirely consistent (Fig. 4E, Supplementary Fig. 4B & 4C).

**Figure 4:**
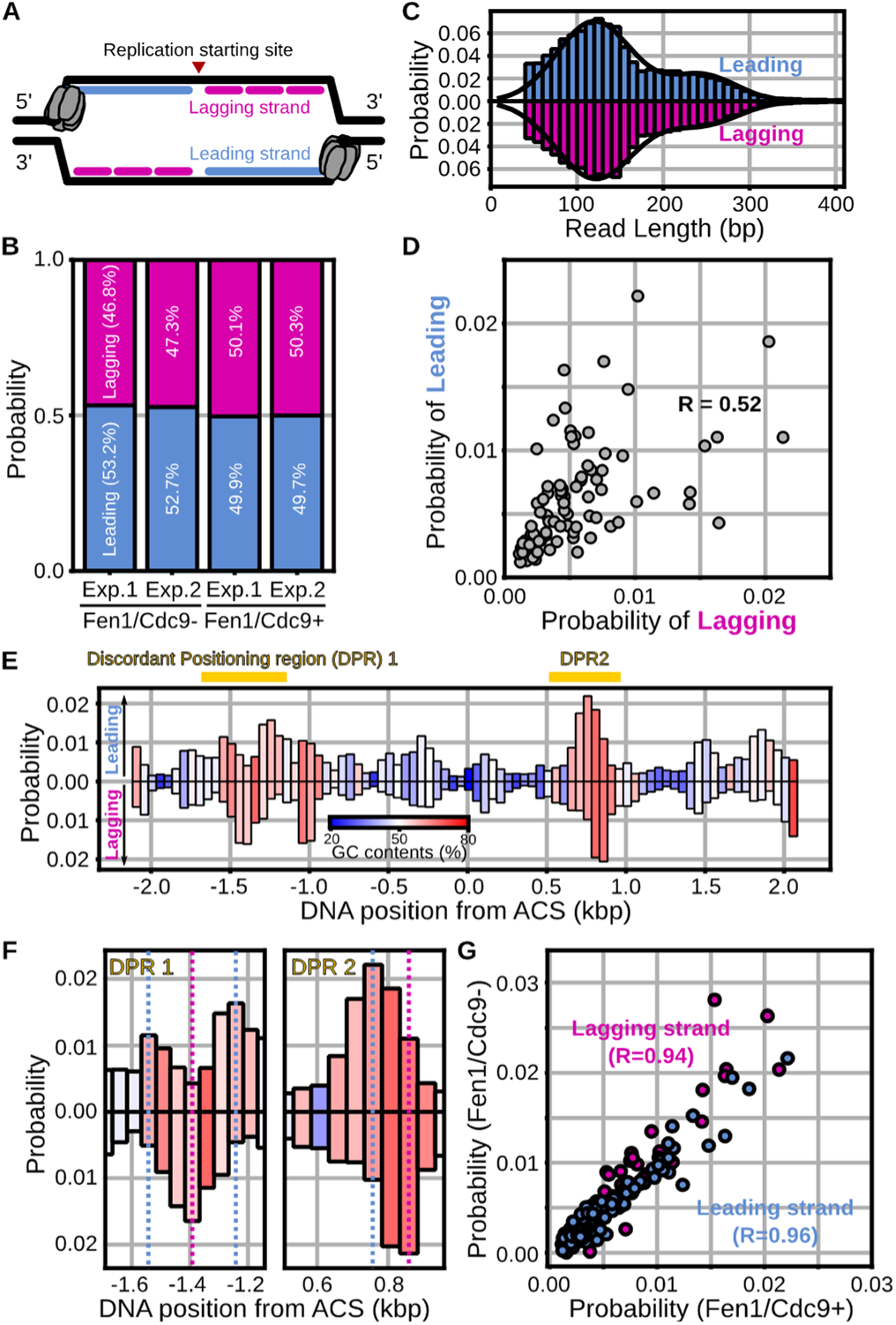
The positions of the histones recycled on the lagging and leading strands. (A) Definition of the leading (cyan) and lagging (magenta) strands. (B) The ratios of the leading and lagging strand fragments in the Fen1/Cdc9− and Fen1/Cdc9+ reactions for two repeated experiments (exp.). (C) The read length distribution of the leading and lagging strand fragments. (D) The correlation plot for the position distributions of recycled histones on the lagging and leading strands. (E) The position distribution of the recycled histones on the leading (top) and lagging (bottom) strands. The colors represent GC contents. (F) Magnified views of discordant positioning regions (DPRs) in (E). The dashed cyan and magenta lines show the peak positions of the recycled histones on the leading and lagging strands, respectively. (G) The correlation plot for the position distributions of recycled histones in the Fen1/Cdc9− and Fen1/Cdc9+ reactions.

We focused on two discordant positioning regions (DPRs) in which the recycled positions were reproducibly inconsistent between the lagging and leading strand (Fig. 4E, 4F & Supplementary Fig. 4D). In the DPR1 (−1.65 to −1.15 kbp away from the ACS site), we found two peaks on the leading strand and one peak on the lagging strand (Fig. 4F & Supplementary Fig. 4D, left). The two peaks on the leading strand were located 100–200 bp away from the single peak on the lagging strand. Moreover, we found that one peak on the lagging strand shifted by ∼100 bp away from the peak on the leading strand in the DPR2 (0.55 to 0.95 kbp away from the ACS site) (Fig. 4F & Supplementary Fig. 4D, right). Since the DNA sequences of the lagging and leading strands are the same, the free energy landscape of nucleosome formation cannot explain the discordant. Instead, the kinetics of replication progression (e.g., speed of DNA polymerases) might be possible factors for the discordant positioning. Interestingly, we noticed that DPRs overlap with GC-rich potential quadruplex-forming sequences where the speed of helicases and DNA polymerases can be slowed down^53^ or uncoupled^54,55^ (Fig. 4E, 4F, Supplementary Fig. 4D, 4E & 4F). Together, we observed the discordance of the recycled position between leading and lagging strands on the GC-rich DNA sequences.

So far, our *in vitro* reconstitution system did not contain Okazaki fragment maturation factors Fen1/Cdc9. To examine the role of the lagging-strand maturation in histone recycling, we applied the Repli-pore-seq pipeline to the reactions with Fen1/Cdc9. The loss of 100–200 bp Okazaki fragment band and appearance of 4 kbp ligated DNA band on the alkaline denaturing gel supported successful maturation of the lagging strand in the presence of Fen1/Cdc9 even on the chromatinized DNA (Supplementary Fig. 5A). Regardless of the maturation, the read length of the lagging and leading strand fragments was not significantly changed (130 ± 43 bp and 131 ± 41 bp, respectively) (Supplementary Fig. 5B). We reproducibly observed that half of the histones were recycled to the leading strand (49.9 % and 49.7 % in the repeated experiments) (Fig. 4B, Supplementary Fig. 5C & 5D), suggesting that the lagging-strand maturation hardly affected the symmetric recycling. Moreover, the position distributions of the lagging and leading strands were highly correlated between the absence and presence of Fen1/Cdc9 (R = 0.94 and 0.96 for the lagging and leading strands, respectively) (Fig. 4G). Together, we found that the lagging-strand maturation factors, Fen1/Cdc9, are not essential for symmetric and faithful recycling, suggesting that maturation may happen after recycling.

### The removal of Pol δ, Ctf4, and Csm3/Tof1 affected histone recycling

As shown in Fig. 1E & 1F, the proteins dispensable for chromatin replication (Pol δ, Ctf4, and Csm3/Tof1) were not strictly required for histone deposition. However, these proteins may affect the recycling symmetry and positions. To investigate the roles of these factors in histone recycling, we performed Repli-pore-seq for the reactions without each factor.

The read length values on the lagging strand fragments in the Pol δ−, Ctf4−, and Csm3/Tof1− reactions were 125 ± 43, 128 ± 49, and 127 ± 42 bp and those of the leading strands were 124 ± 40, 118 ± 45, and 124 ± 35 bp (Supplementary Fig. 6A). The removal of each factor did not significantly alter the read length, consistent with the gel-electrophoresis-based assays (Fig. 1E & 1F). The fractions ranging from 40 to 120 bp increased after replication in the same way as the presence of all factors (Fig. 3B, 4C & Supplementary Fig. 6A). Therefore, the results suggested that some nucleosomes lost one or two H2A/H2B dimers to form hexasomes or tetrasomes after histone recycling without Pol δ, Ctf4, or Csm3/Tof1.

Next, we investigated the effects of Pol δ, Ctf4, and Csm3/Tof1 on the recycling symmetry. Of the genuinely biotinylated DNA fragments, 61.9%, 55.0%, and 52.9% were the leading strand fragments in the Pol δ−, Ctf4−, and Csm3/Tof1− reactions, respectively (Fig. 5A). We also observed the statistically significant leading-strand bias in the repeated experiments of Pol δ− (62.0%) and the slight or no bias in the repeated experiments of Ctf4− (55.0%) or Csm3/Tof1− (53.0%) (Fig. 5A). For both the lagging and leading strand fragments, the position distributions of the Pol δ−, Ctf4−, and Csm3/Tof1− reactions highly correlated with those in the presence of them (Fig. 5B). Moreover, the comparison of the deposition probabilities to each position between the Pol δ− and Pol δ+ reactions revealed the global decrease (Fig. 5B), consistent with the compromised lagging-strand synthesis upon Pol δ removal (Fig. 1B & 1C). These results indicated that Pol δ contributes to the recycling symmetry. In contrast, the effects of Ctf4 and Csm3/Tof1 were miniscule.

**Figure 5:**
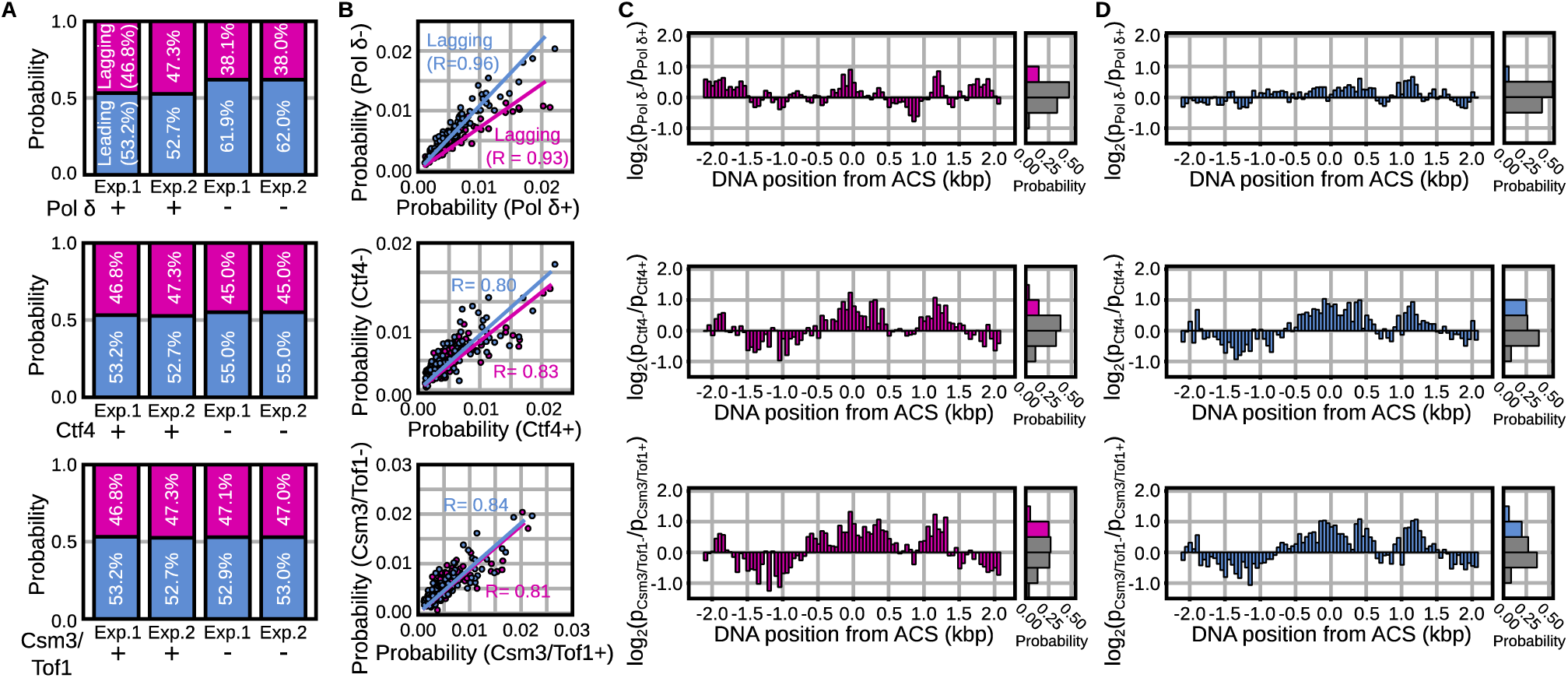
Effects of removing Pol δ, Ctf4, and Csm3/Tof1 on histone recycling. (A) The ratios of the leading and lagging strand fragments in the Pol δ− and Pol δ+ reactions (top), the Ctf4− and Ctf4+ reactions (middle), and the Csm3/Tof1− and Csm3/Tof1+ reactions (bottom) for two repeated experiments (exp.). (B) The correlation plot for the position distributions of recycled histones on the lagging and leading strands in the Pol δ− and Pol δ+ reactions (top), the Ctf4− and Ctf4+ reactions (middle), and the Csm3/Tof1− and Csm3/Tof1+ reactions (bottom). (C & D) The logarithmic fold change (log_2_FC) of the position distributions of recycled histones on the lagging (C) and leading (D) strands in the Pol δ− and Pol δ+ reactions (top), the Ctf4− and Ctf4+ reactions (middle), and the Csm3/Tof1− and Csm3/Tof1+ reactions (bottom). In the histograms, the bars were colored gray if the log_2_FC values was <0.5; otherwise, they were colored magenta (C) or cyan (D).

We also investigated the effects of Pol δ, Ctf4, and Csm3/Tof1 on the recycling positions. We normalized the position distribution of the lagging and leading strand fragments by their ratios of reads Then, we calculated the fold change (FC) of the normalized distributions before and after the removal. In the Pol δ−, Ctf4− and Csm3/Tof1− reaction, 14%, 18%, and 30% positions of the lagging strand fragments and 5%, 26%, and 26% positions of the leading strand fragments showed the log_2_FC value above 0.5, respectively (Fig. 5C & 5D). Upon the removal of each factor, the log_2_FC values increased over the positions within −0.5 to 0.5 kbp and 1.0 to 1.5 kbp from the ACS site for both the lagging and leading strand fragments (Fig. 5C & 5D). These regions correspond to the DNA sequences energetically unfavorable for forming nucleosomes (Fig. 3E). Therefore, the results indicated that the removal of Ctf4 or Csm3/Tof1 altered the histone positions to the energetically unstable ones (Fig. 5C & 5D). Together, the removal of Ctf4 or Csm3/Tof1 caused alterations in the recycling positions.

## DISCUSSION

In this study, we successfully reconstituted histone recycling using ∼30 purified proteins and analyzed the outcomes with a newly developed method called Repli-pore-seq, which combines nanopore sequencing and deep-learning-based classification. Our results revealed that histones are symmetrically recycled onto the lagging and leading DNA strands, forming tetrasomes or hexasomes on sequences favoring nucleosome assembly. Notably, we observed positional discordance of recycled histones between the two replicated strands in GC-rich regions despite identical DNA sequences. Additionally, we demonstrated that the lagging-strand maturation factors, Fen1/Cdc9, are not required for symmetric recycling, implying that maturation occurs after recycling. While gel-electrophoresis-based assays showed that Pol δ, Ctf4, and Csm3/Tof1 are not essential for histone deposition, the absence of Pol δ disrupted recycling symmetry, and removing Ctf4 or Csm3/Tof1 altered recycling positions.

Okazaki fragment (OF) maturation of the lagging strand is essential to complete DNA synthesis. Pol δ, which extends an OF, invades and displaces the 5’-end of the adjacent OF to produce a flap structure. Fen1 removes the flap, and Cdc9 ligates the two OFs. Previous studies have demonstrated that the junction of two OFs is enriched around the nucleosome dyad and that the distribution of the junction can be altered by impairing chromatin assembly or Pol δ processivity^56–58^. Moreover, Fen1/Cdc9 can catalyze the flap excision and ligation on nucleosome substrates^59,60^. The current study revealed that the Okazaki fragment maturation does not alter the oligomeric state of the recycled histones and recycling symmetry and positions. These results suggested that histone recycling occurs before maturation, which can proceed without disrupting epigenetic information encoded in histones.

Recent studies identified multiple proteins [Mcm2, Pol α, Ctf4, Mrc1, Tof1, Pol32 (Pol δ), FACT, PCNA, and Dpb3/4], which cooperatively work to ensure symmetric histone recycling^16–26^. In particular, recent eSPAN analyses have revealed that Ctf4 deletion (*ctf4Δ*)^18^ and Tof1 mutations in residues interacting with the N-terminal tail of Mcm2 (*tof1-3A*)^21^ disrupted histone recycling to the lagging strand. In the current study, however, the effects of Ctf4 or Csm3/Tof1 removal on the recycling symmetry were minuscule. This discrepancy suggests that additional unknown factors dictate the recycling symmetry in cooperation with Ctf4 and Csm3/Tof1, which are absent in our reconstitution. Future studies focusing on these missing components will be crucial to fully elucidate the interaction network governing histone recycling during DNA replication.

Faithful epigenetic inheritance is essential for the maintenance of heterochromatin through generations^13,22,23^. Histone recycling is the first step of heterochromatin maintenance, followed by incorporating newly synthesized histones, restoring epigenetic marks, and re-organizing chromatin structure. The subsequent processes of chromatin maturation require additional proteins such as histone chaperones (CAF1 and Asf1), histone-modifying enzymes, and proteins to organize chromatin (HP1, linker histones, and SMC-proteins)^4–7^. Combination of *in vitro* chromatin replication reconstitution with Repli-pore-seq allowed adding proteins to or removing those from the system and analyzing the chromatin structure. Therefore, the pipeline developed here should be useful for studying molecular mechanisms of chromatin maturation.

The analyses of the recycling positions revealed discordance between the lagging and leading strands at GC-rich and G-quadruplex-prone regions. Previous studies showed the slowing down of DNA polymerases and helicase/polymerase uncoupling in GC-rich regions^53–55^. Moreover, recent studies have suggested that recycling symmetry is modulated by the rate balance of the lagging and leading DNA synthesis^42,61^. These results naturally lead to the hypothesis that the discordance of the recycling positions reflects the uncoupling of the lagging and leading strand synthesis. Techniques to directly visualize the dynamics of replication components (e.g., single-molecule fluorescence imaging and molecular dynamics simulations) can evaluate the hypothesis in the future.

The Repli-pore-seq pipeline combines the reconstituted eukaryotic DNA replication and computational “purification” of the replicated DNA using deep learning-based classification. *In vitro* reconstitution studies have an advantage in studying protein mutation and depletion, which have lethal effects on cells. On the other hand, the reconstitution has a limitation regarding low reaction scale and efficiency. Even a single biochemical purification step after the reconstitution reaction sometimes has a detrimental effect on the yield of the subsequent analysis. Our computational purification procedure allows bypassing the immunoprecipitation, which resulted in the loss of replicated DNA in our initial trial. Therefore, the Repli-pore-seq pipeline expands the scope of sequencing-based analysis to the *in vitro* reconstitution system.

## METHODS

### Material preparations

The 4.1 kbp plasmid, pARS1-4.1, was used as the DNA substrate of *in vitro* replication assays. The DNA fragments to make training data for the deep-learning model were amplified with primers 5’-CCATTTGTCTCCACACCTCC-3’ and 5’-CCATTTGTCTCCACACCTCC-3’ by PCR. The biotinylated fragments were amplified in the reaction containing 200 μM each of dGTP, dCTP, and dATP, and 100 μM each of dTTP and biotin-dUTP. The budding yeast histones (H3, H4, H2A, and H2B) and replication proteins (ORC, Cdc6, Mcm2-7/Cdt1, DDK, S-CDK, Sld2, Sld3/7, Dpb11, Cdc45, GINS, PCNA, RFC, Mcm10, RPA, DNA polymerase α, DNA polymerase δ, DNA polymerase ε, Ctf4, Csm3/Tof1, Mrc1, and Fen1/Cdc9) were purified as described previously^41^.

### *In vitro* reconstitution of chromatin replication

The nucleosomes were reconstituted by mixing the pARS1-4.1 plasmid DNA (4 nM) with histone octamers (400 nM), ISW1a (17.3 nM), Nap1 (3 μM), and ATP as described previously^41^. After the nucleosome assembly, we exchanged the buffer and removed residual proteins using MicroSpin S-400 HR column (Cytiva). Then, we assembled MCM double hexamer onto the chromatinized DNA substrates by adding 20 nM ORC, 50 nM Cdc6, and 60 nM Mcm2-7/Cdt1 in the MCM loading buffer [25 mM HEPES-KOH (pH 7.5), 100 mM KOAc, 1 mM DTT, 5 mM ATP, 7.5 mM Mg(OAc)_2_, 5% glycerol, 0.01% (w/v) NP-40, and 0.1 mg/mL BSA]. After 15 minutes incubation at 30 °C, 40 nM DDK was added to the reaction and further incubated for 15 minutes. Following the MCM loading, the reaction was diluted three-fold in the replication buffer [25 mM HEPES-KOH (pH 7.5), 100 mM KOAc, 1 mM DTT, 5 mM ATP, 7.5 mM Mg(OAc)_2_, 5% glycerol, 0.01 % (w/v) NP-40, 0.1 mg/mL BSA, 0.1 mM CTP, 0.1 mM GTP, 0.1 mM TTP, 40 μM dATP, 40 μM dCTP, 40 μM dGTP, 25 μM dTTP, and 15 μM biotin-dUTP], and mixed with 5 nM S-CDK, 30 nM Sld2, 20 nM Sld3/7, 30 nM Dpb11, 40 nM Cdc45, 12.5 nM GINS, 20 nM Csm3/Tof1, 20 nM Ctf4, 10 nM Mrc1, 20 nM RFC, 20 nM PCNA, 100 nM RPA, 5 nM Pol α, 10 nM Pol δ, and 20 nM Pol ε, 6.7 nM ORC, 16.7 nM Cdc6, and 20 nM Mcm2-7/Cdt1, 13.3 nM DDK. We started the replication reaction by adding a histone chaperone FACT (40 nM FACT and 200 nM Nhp6A) and a firing factor Mcm10 (5 nM) to the reaction mixture and incubated at 30℃ for 20 minutes. After the replication reactions, we adjusted the salt concentration to 50 mM NaCl and 2 mM CaCl_2_ and digested the replication products by adding 1 U/μL of Micrococcal nuclease (MNase) and placing it at 30°C for 30 minutes. Then, the reaction was mixed with 15 mM EDTA, 0.1% (w/v) SDS, and 0.5 mg/mL protease K and incubated at 37 °C for 20 minutes to stop the reaction.

For native polyacrylamide gel electrophoresis, the samples were mixed with the 6× loading buffer [15% (w/v) ficol, 10 mM HEPES-KOH (pH 7.5), 0.05% orange G] and were applied to 7.5% polyacrylamide gel in EzRun TG buffer (ATTO) at 21 mA at room temperature for 65 minutes. The gel was soaked in 0.5× TBE for 5 minutes, and the DNA fragments were transferred to the Zeta-Probe membrane (Bio-Rad) using a wet-transfer blotter at 80 V at 4°C for 50 minutes. The transferred membrane was crosslinked using a UV illuminator and soaked in the blocking solution (Cytiva) at room temperature for 20 minutes. The membrane was incubated with Dylight680-conjugated streptavidin in the TBS-T buffer [TBS containing 0.1% (w/v) Tween 20 and 0.01% (w/v) SDS] for 30 minutes, followed by washing the membrane three times with the TBS-T buffer. Gel images were captured by ChemiDoc Touch imager (Bio-Rad).

For denaturing agarose gel electrophoresis, the sample was mixed with 50 mM EDTA and the 6× loading buffer [0.3 M NaOH, 6 mM EDTA, 36% (w/v) Glycerol, 0.25% (w/v) orange G] and were applied to 0.8% agarose gel in the alkaline buffer (50 mM NaOH, 1 mM EDTA) at 3.6 V/cm at 4°C for 2 hours. The gel was dipped in 0.5× TBE for 10 minutes, and the DNA fragments were transferred to the nylon membrane (Hybond-N+, Cytiva) using a wet-transfer blotter at 80 V at 4°C for 50 minutes. The biotinylated DNA fragments were detected as described above.

### Nanopore sequencing

For nanopore sequencing, the DNA fragments in the samples digested by MNase were purified in sterilized water using the silica membrane spin column (Qiagen). We repaired the purified DNA fragments using the NEBNext® Companion Module for Oxford Nanopore Technologies® Ligation Sequencing (New England Biolabs) and ligated the adapter proteins to the repaired DNA fragments using the Ligation Sequencing Kit V14 (SQK-LSK 114, Oxford Nanopore Technologies®). We loaded the DNA library onto the MinION R10.4.1 flowcell (FLO-MIN 114, Oxford Nanopore Technologies®) and ran the sequencing on MinKNOW (Oxford Nanopore Technologies®). The current signals were translated into nucleotide sequences using the Guppy software (Oxford Nanopore Technologies®).

### Deep-learning model training

We used the current signals of the PCR-amplified biotinylated and unbiotinylated DNA substrates as training data for our deep-learning model. We segmented the current signals based on a stepwise change, calculated each segment’s mean, standard deviation, and duration time, and converted the signals into an array of segments. When the number of segments in one array exceeded 600, the 200– 600 subset segments were randomly extracted. Then, we padded the arrays to the size of 3 × 600. The segment arrays derived from unbiotinylated and biotinylated DNA were labeled 0 and 1, respectively. We kept running the sequencing of unbiotinylated and biotinylated DNA so that their sequence coverages were to be (4.8 ± 0.8) × 10^4^ and (6.2 ± 1.3) × 10^4^, respectively. All the obtained current signals were converted into 538,705 arrays each of unbiotinylated- and biotinylated-DNA, and randomly split the total (2 × 538,705) arrays into 969,669 and 107,741 arrays for training and validation data, respectively.

Each array of 3 × 600 size was fed into the depth-wise and point-wise convolution, batch normalization, and ReLU activation (DSC-PSC-BN-ReLU) layers. The output tensors were fed into 8 repeats of the residual block followed by LSTM with 128 units and the max pooling layer. Each residual block contained 4 sets of DSC-PSC-BN-ReLU followed by one DSC-PSC-BN. The output from the network was fully connected and activated by sigmoid functions. We used focal cross entropy as a loss function. The model was trained using Adam optimizer with a learning rate of 0.001 for 1000 epochs with a batch size of 1024. The output values from the final layer below 0.5 were regarded as ‘unbiotinylated’ while those of 0.5 and above were considered ‘biotinylated.’

### Correction for misclassified reads

In the Mcm10− reactions, ∼0.6% of the reads were misclassified as ‘biotinylated’ (Supplementary Fig. 2C). Thus, the biotinylated reads should be a mixture of genuinely biotinylated reads and misclassified unbiotinylated reads. To obtain an ensemble of genuinely biotinylated reads, we need to subtract the contribution of the misclassified reads from an ensemble of biotinylated reads in the Mcm10+ reactions.

We define the number of total reads, biotinylated reads, unbiotinylated reads, misclassified biotinylated reads, genuinely biotinylated reads, and genuinely unbiotinylated reads as *N*, *N_Positive_*, *N_Negative_*, *N_False-Positive_*, *N_Biotinylated_*, and *N_Unbiotinylated_*, respectively. Then, we have the following relationships.

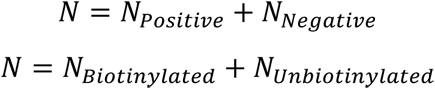

Based on the assumption that biotinylated reads are a mixture of truly biotinylated reads and misclassified unbiotinylated reads, the following relationship holds.

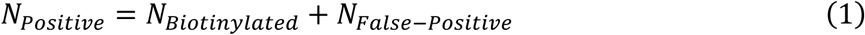

The ratios of ‘biotinylated’ reads, misclassified unbiotinylated reads, and truly biotinylated reads to total reads (*r_Positive_*, *r_False-Positive_*, and *r_Biotinylated_*) are given by,

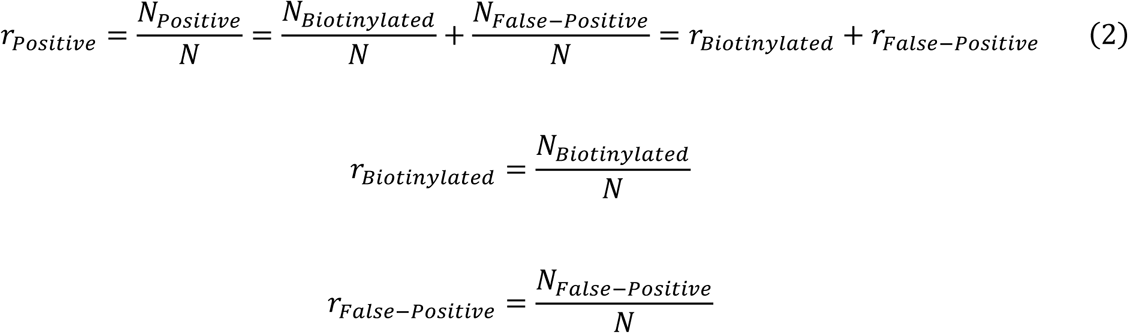

Here, we assume that the classifier mistakenly classifies genuinely unbiotinylated reads as biotinylated reads at a ratio α. Then, the number and ratio of misclassified unbiotinylated reads are given by,

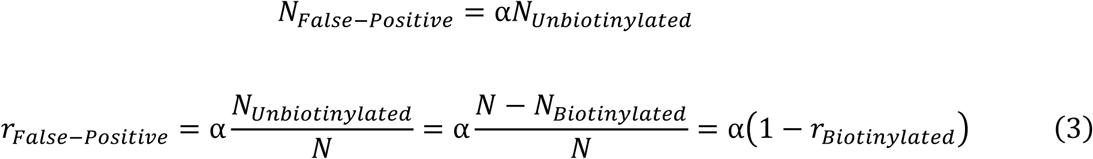

From equation 2 and 3, we can derive the following relationships:

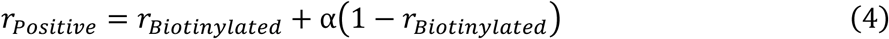

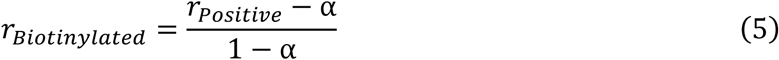

In the Mcm10− reaction, all reads should be unbiotinylated ones. Then, r_Biotinylated_ is equal to 0 in equation 5, and the following relationship holds.

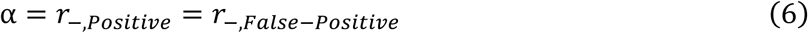

Next, we estimated an ensemble of truly biotinylated reads. In this study, we obtained the position distributions of the DNA fragments classified to be biotinylated in the Mcm10+ reaction. As shown in Supplementary Fig. 2E, we also obtained the position distributions of the DNA fragments misclassified to be biotinylated in the Mcm10− reaction. We need to subtract this misclassification bias from the position distributions of the biotinylated reads. Here, we define the position probabilities of the total, genuinely, and misclassified biotinylated reads at position *x* as *p_Positive_*(*x*), *p_Biotinylated_*(*x*), and *β*(*x*), respectively. Then, the numbers of total [*n_Positive_*(*x*)], genuinely [*n_Biotinylated_*(*x*)], and misclassified [*b*(*x*)] biotinylated reads at position *x* are represented as

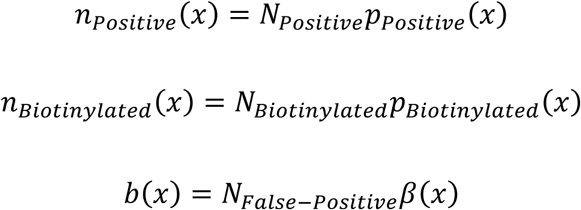

Here, the following relationships hold as in equation 1.

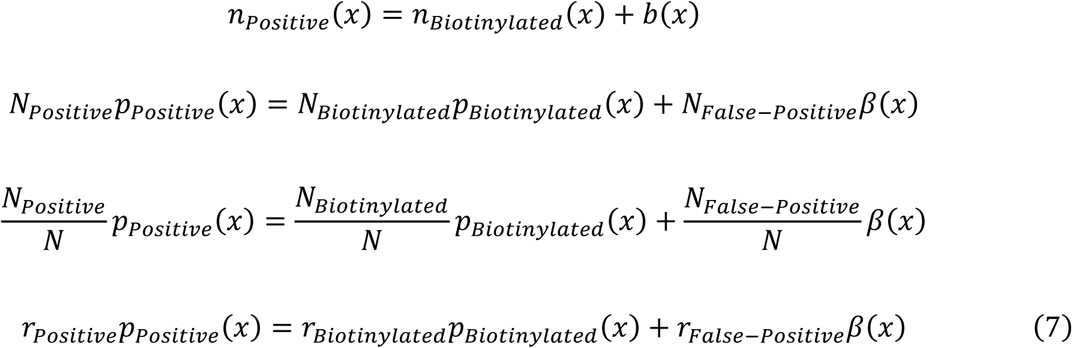

From equations 3 and 7, we can derive the following equations.

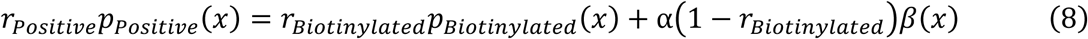

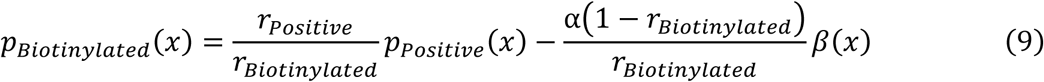

From equations 5 and 9, we can derive the following equation.

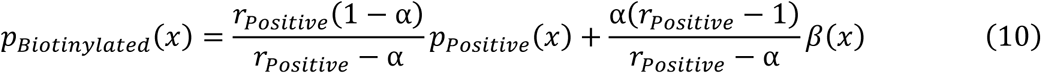

Again, in the Mcm10− reaction, the ratio of truly biotinylated reads should be 0. Thus, from equations 6 and 8, we can derive the following relationships:

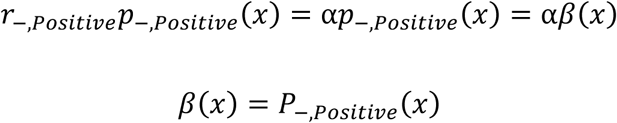

Therefore, *β*(*x*) can be calculated from the position distributions of the DNA fragments misclassified to be biotinylated in the Mcm10− reaction. Then, we can subtract the contribution of the misclassified reads from an ensemble of biotinylated reads in the Mcm10+ reactions using equation 10.

### Prediction of potential quadruplex-forming sequence

The quadruplex-forming sequences (PQS) were predicted using pqsfinder^62^. The PQS was searched as four guanine runs (G-runs). Each G-run was defined as a stretch of 2 to 20 guanines, with one insertion of a stretch of 0–9 other nucleotides allowed. The score function to evaluate the PQS site is based on the reward for G-tetrad stacking and on the penalty for mismatches and the insertion in G-runs. The parameters were calibrated by maximizing the correlation between the signal from G4-seq dataset^63^ and the score.

## AKNOWLEDGEMENTS

We thank Prof. Shoji Takada and the laboratory members of the theoretical biophysics laboratory at Kyoto University for discussions and assistance throughout this work. Also, we thank Prof. Hitoshi Kurumizaka and his laboratory members for helping us to purify histones and reconstitute nucleosomes. This work was supported by the Grant-in-Aid for Transformative Research Areas (24H00883; to T.T.), the grant from the Kyoto University Foundation (to T.T.), the grant from the Takeda Science Foundation (to T.T.), the grant from the Shimazu Science Foundation (to T.T.), the grant from the Inamori Foundation (to T.T.), the Grant-in-Aid for Japan Society for the Promotion of Science Fellows (22J21003; to F.N.), and the grant from the Ginpuu Foundation (to F.N. and T.T.).

## DATA AVAILABILITY

The data that support the findings of this study are available within the article and Supplementary Information files. The plasmid DNA sequence for pARS1-4.1 and the codes for the deep-learning based classifier are available at https://github.com/TerakawaLab/2024_Nagae.

## AUTHOR CONTRIBUTIONS

F.N., Y.M., and T.T. designed the project. F.N. performed the nanopore sequencing, the gel-electrophoresis-based assays and their analyses, S.E. and Y.M. performed the *in vitro* chromatin replication assays, the gel-electrophoresis-based assays, and their analyses. F.N., Y.M., and T.T. co-wrote the manuscript.

## SUPPLEMENTARY INFORMATION

**Supplementary Figure 1:**
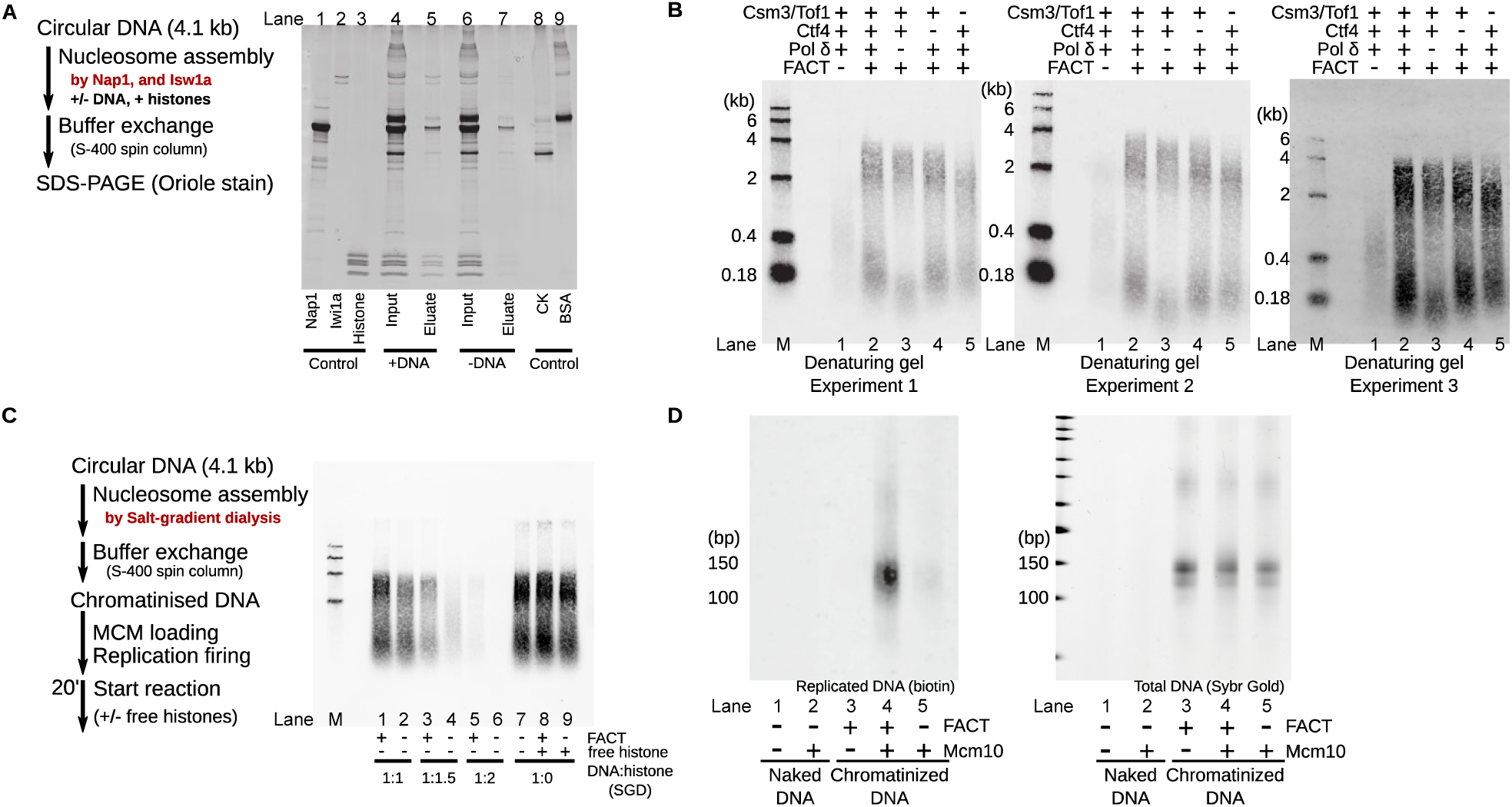
*In vitro* reconstitutions of chromatin replication. (A) The SDS polyacrylamide gel image of denatured proteins in the nucleosome reconstitution reactions before (input) and after (eluate) buffer exchange by the S-400 spin column. (B) The alkaline denaturing agarose gel images of replicated DNA in three repeated experiments using the NI-chromatin as substrates. ‘M’ denotes a marker. (C) The alkaline denaturing agarose gel images of replicated DNA using the SGD-chromatin as substrates. ‘M’ denotes a marker. (D) The native polyacrylamide gel images of total (right) and replicated (left) DNA after MNase digestion.

**Supplementary Figure 2:**
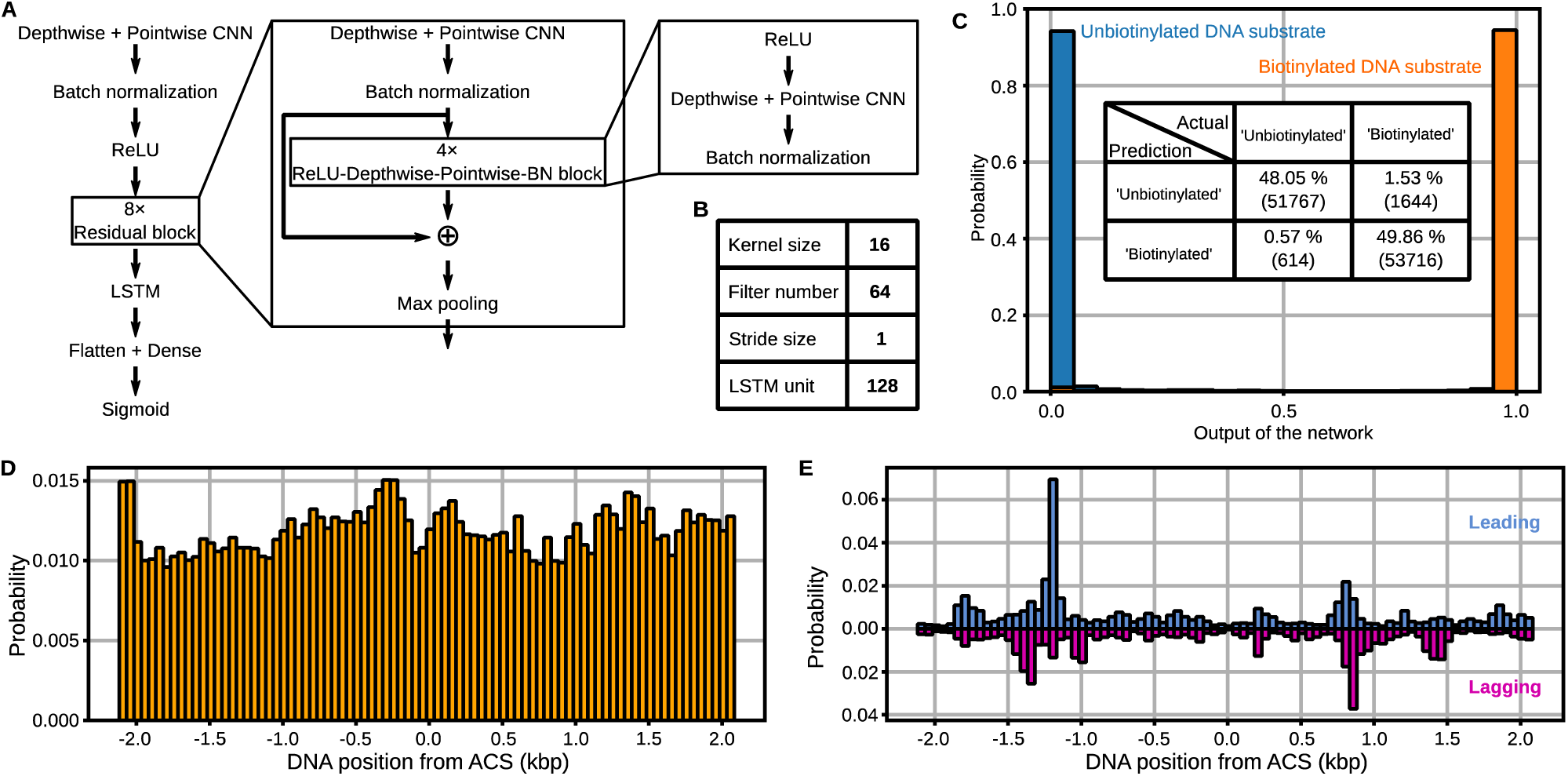
The deep-learning model. (A) The architecture of the neural network (B) The parameters of the convolution and LSTM layers. (C) The probability distributions of the model outputs of the validation datasets. The inset table is the confusion matrix. (D) The position distributions of the DNA fragments from naked plasmid sheared by sonication. (E) The position distribution of the histones on the lagging and leading strands in the Mcm10− reaction.

**Supplementary Figure 3:**
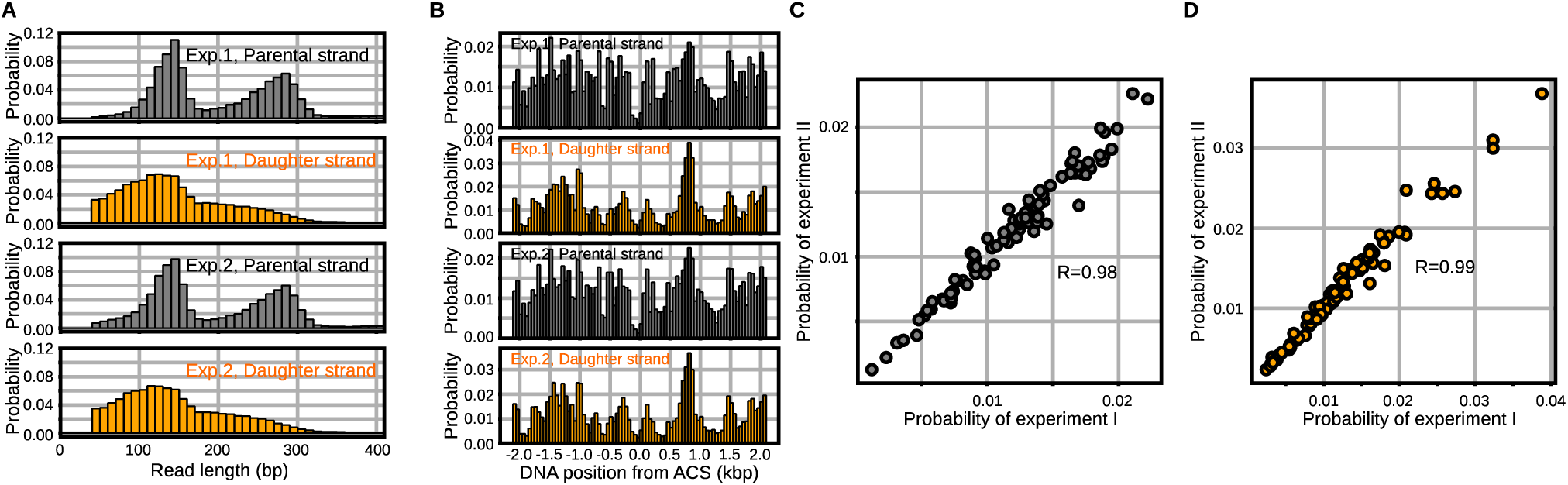
Repli-pore-seq of the reconstituted chromatin replication. (A) The read length distributions of parental DNA fragments (top, gray) and daughter DNA fragments (bottom, orange) for two repeated experiments (exp.). (B) The position distributions of histones on the parental DNA (top; gray) and the daughter DNA (bottom; orange) for two repeated experiments (exp.). (C & D) The correlation plot for the position distributions of histones on the parental (C) and daughter (D) DNA from two repeated experiments in (B).

**Supplementary Figure 4:**
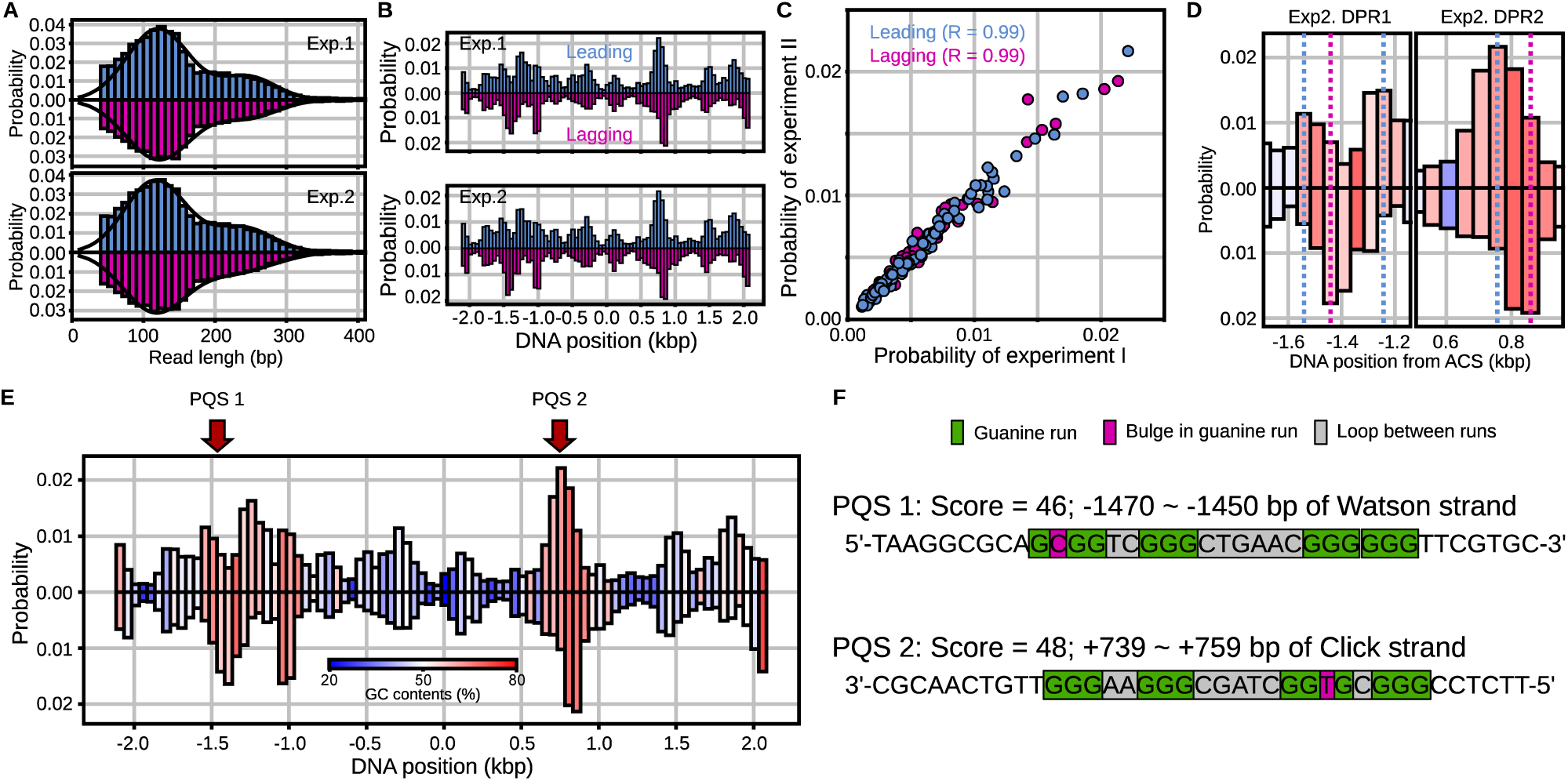
The positions of the histones recycled on the lagging and leading strands. (A) The read length distribution of the leading (cyan) and lagging (magenta) strand fragments for two repeated experiments (exp.). (B) The position distributions of recycled histones on the leading (top) and lagging (bottom) strands for two repeated experiments (exp.). (C) The correlation plot for the position distributions of recycled histones on the lagging and leading strands for two repeated experiments. (D) Magnified views of discordant positioning regions (DPRs) in Fig. 4E but for the second experiment. (E) The position distribution of the recycled histones on the leading (top) and lagging (bottom) strands. The colors represent GC contents. (F) Sequences of the predicted PQS sites.

**Supplementary Figure 5:**
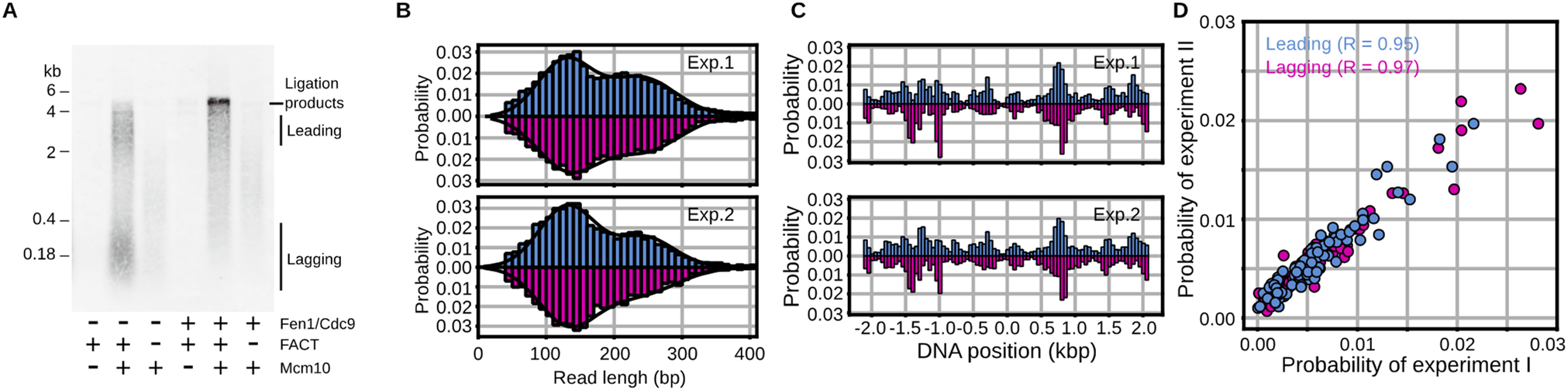
The positions of the histones recycled on the lagging and leading strands in the Fen1/Cdc9+ reactions. (A) The read length distribution of the leading (cyan) and lagging (magenta) strand fragments for two repeated experiments (exp.). (C) The position distributions of recycled histones on the leading (top) and lagging (bottom) strands for two repeated experiments (exp.). (D) The correlation plot for the position distributions of recycled histones on the lagging and leading strands for two repeated experiments.

**Supplementary Figure 6:**
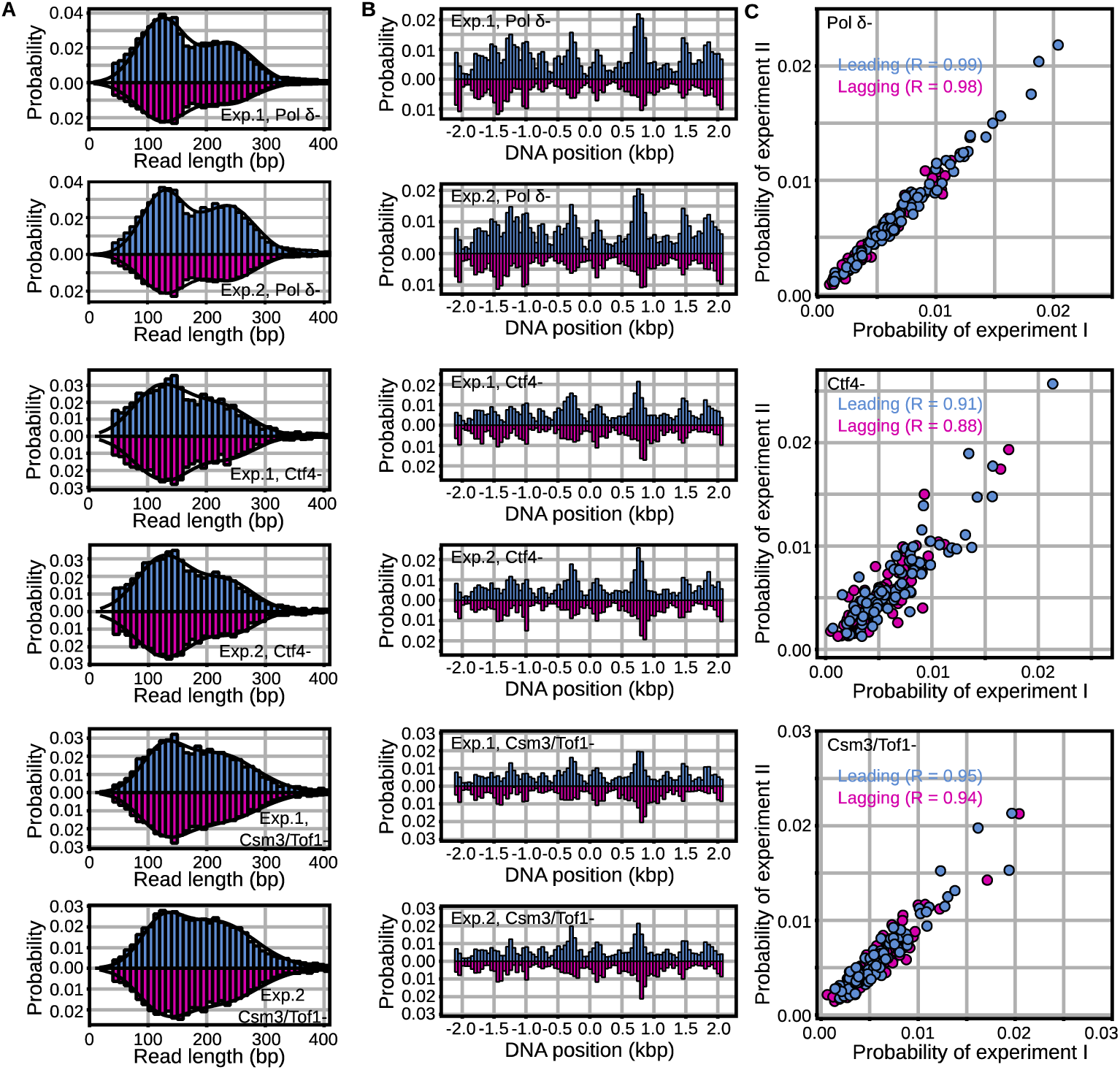
Effects of removing Pol δ, Ctf4, and Csm3/Tof1 on histone recycling. (A) The read length distribution of the leading (cyan) and lagging (magenta) strand fragments in the Pol δ− and Pol δ+ reactions (top), the Ctf4− and Ctf4+ reactions (middle), and the Csm3/Tof1− and Csm3/Tof1+ reactions (bottom) for two repeated experiments (exp.). (B) The position distributions of recycled histones on the leading (top) and lagging (bottom) strands in the Pol δ− and Pol δ+ reactions (top), the Ctf4− and Ctf4+ reactions (middle), and the Csm3/Tof1− and Csm3/Tof1+ reactions (bottom) for two repeated experiments (exp.). (C) The correlation plot for the position distributions of recycled histones on the lagging and leading strands in the Pol δ− and Pol δ+ reactions (top), the Ctf4− and Ctf4+ reactions (middle), and the Csm3/Tof1− and Csm3/Tof1+ reactions (bottom) for two repeated experiments (exp.).

